# Glyceraldehyde-3-Phosphate Dehydrogenase and Seven in Absentia Homolog -1 Interactions in Mammalian Cells Visualized Using Proximity Ligation Assay

**DOI:** 10.1101/090068

**Authors:** Harkewal Singh, Christopher Melm

## Abstract

Proteins seldom function in isolation and thus protein-protein interactions are critical in understanding the molecular basis of diseases and health (*1*, *2*). There are several well established techniques that are used to investigate protein-protein interactions(*3*). Most of the methods require some form of genetic modification of the target protein and thus always adds extra steps. However, Proximity Ligation Assay(*4*-*6*) (PLA) *aka* Duolink® is one such method that requires no genetic modification of the target protein and probes protein-protein interactions in fixed live cells and tissues. Briefly, PLA requires the use of primary antibodies specific to the proteins of interest. Once the sample (fixed cells or tissues) is incubated with species specific primary antibodies, secondary antibodies that are conjugated with oligonucleotides (also known as PLUS and MINUS probes respectively) and connecter oligonucleotides are added. This complex is ligated if the two PLUS and MINUS probes are within 40nm of each other. The resulting nucleic acid is amplified using rolling circle amplification and then probed with appropriate fluorescent probes. If the two proteins are interacting, one could visualize the interaction as a single red foci (for example Far Red Detection) using fluorescent microscopy. Here, we used PLA to probe protein-protein interactions between Glyceraldehyde-3-phosphate dehydrogenase (GAPDH) and seven in absentia homolog -1 (Siah-1) – an E3 ubiquitin ligase. We first use PLA to show that GAPDH and Siah-1 proteins exist endogenously in the cytosol of multiple mammalian cell lines. Our data suggest the use of DU145 and T98G cell lines to show translocation of the GAPDH-Siah- 1 complex. Next, we used common nitrosylation agents(*7*, *8*) (S-nitrosoglutathione-GSNO and S-Nitroso-N-acetyl-DL-pencillamine–SNAP) in different concentrations and observed that GAPDH and Siah-1 interact presumably due to the nitrosylation of the former, which is consistent with previous studies(*9*, *10*). Interestingly, no interactions were observed between the two proteins in the absence of GSNO or SNAP indicating that nitrosylation might be critical for GAPDH-Siah1 interactions. Our results suggest that GAPDH-Siah-1 interactions originate in the cytosol and migrate to the nucleus under the conditions tested. We quantify the PLA signal using Duolink® Image Tool and observe a clear enhancement of GAPDH-Siah-1 PLA signal upon treating the cells with GSNO or SNAP. Next, we used R-(-)-Deprenyl (deprenyl), a known inhibitor of GAPDH^4^, and show that it abrogates GAPDH-Siah-1 PLA complex under the conditions tested. Finally, our data suggest that PLA can detect and quantify the GAPDH-Siah1 complex; a well-known protein-protein interaction implicated in neurodegeneration(*9*-*11*) and thus could be a method of choice for similar applications.

## Introduction

Glyceraldehyde-3-phosphate dehydrogenase (GAPDH) is a cytosolic homotetrameric protein (Figure 1a) with key glycolytic functions and has often been viewed as a housekeeping gene. Interestingly, GAPDH that lacks a nuclear localization signal (NLS) has been shown to have functions in nuclear events such as gene transcription, RNA transport, DNA replication and cell signaling (*12-14*). Furthermore, it has been (*9, 15, 16*) suggested that activation of neuronal nitric oxide synthase (nNOS) leads to S –nitrosylation of GAPDH, abolishing its catalytic activity while conferring the ability to bind seven in absentia homolog -1 (Siah-1) – an E3 ubiquitin ligase (Figure 1b) that contains an NLS. The nuclear localization signal of Siah-1 mediates GAPDH translocation to the nucleus as GAPDH-Siah-1 complex. In the nucleus, GAPDH is proposed to bind protein acetyltransferase (p300/CBP) and increases p300/CBP’s acetylation efficiency(*17*). Curiously, p53 in the nucleus undergoes p300/CBP mediated acetylation as a response to cell insult such as nitrosylation, which subsequently increases cytotoxicity and causes apoptotic cell death (*18*). Discovery (*17*) that NO S-nitrosylation of GAPDH leads to GAPDH-Siah-1 interaction and subsequent GAPDH translocation to the nucleus addresses the role of GAPDH in apoptosis. Several studies reported GAPDH nucleus accumulation under apoptotic conditions despite the fact that GAPDH lacks NLS. It has been shown that GAPDH binds to Siah-1 following S- nitrosylation of the catalytic cysteine of GAPDH, stabilizes otherwise unstable Siah-1, and GAPDH-Siah-1 complex translocates to the nucleus (Figure 2) resulting in degradation of the nuclear targets of Siah-1 and subsequent enhancement of apoptosis (*9*). Thus, GAPDH is not only recognized as NO stress sensor but also GAPDH - Siah-1 complex is implicated in various diseases such as Alzheimer’s disease (AD), Parkinson’s disease(*19*), cerebral ischemia–reperfusion injury(*20*), and acute lung injury(*21*). Interestingly, compounds (deprenyl and CGP-3466 [dibenzo-(b,f)oxepin-10-ylmethyl-methyl-prop-2-ynylamine]) that inhibit GAPDH have been shown to act as potential drug candidates for PD and other neurodegenerative disorders, likely by disrupting the GAPDH - Siah-1 complex (*11*). The diverse role of GAPDH-Siah-1 protein-protein complex in human health and disease makes this protein-protein association an interesting target for finding anti-apoptotic and anti-neurodegenerative compounds(*22*).

**Figure 1.**
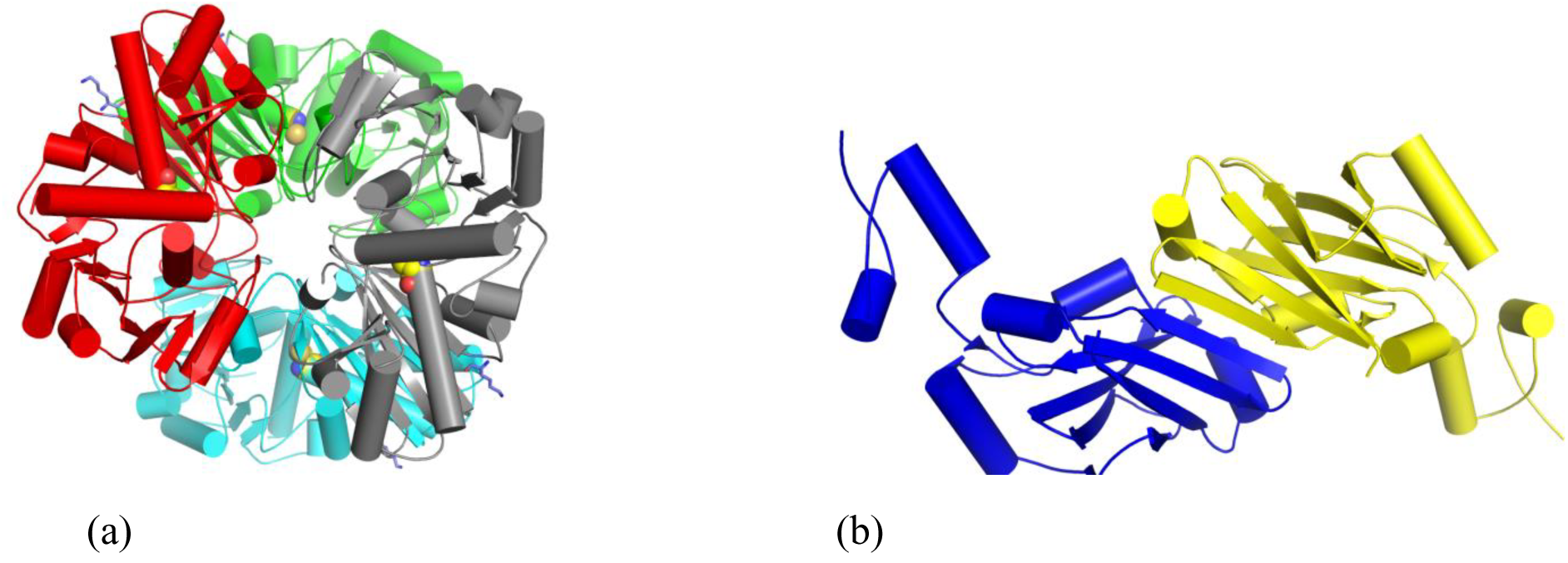
Cartoon representation of a)- GAPDH tetramer5 (PDB code 1U8F) and b)- Siah-16 (PDB code 1K2F).

**Figure 2.**
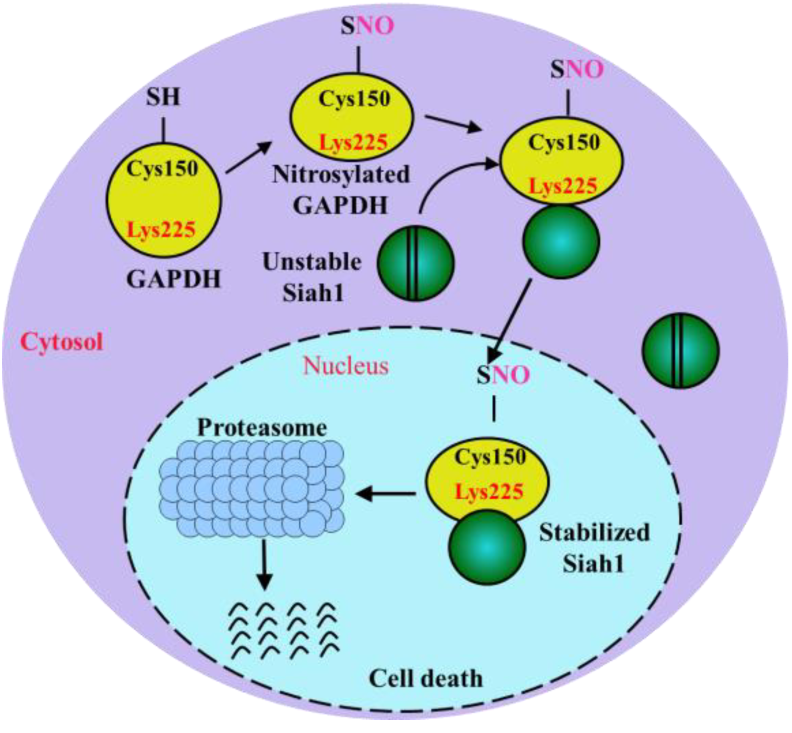
Schematic diagram of GAPDH-Siah1 cascade. Figure shows the proposed nitrosylation of GAPDH, GAPDH association with Siah-1 and GAPDH-Siah1 complex translocation. Figure adapted from Hara et al.

Here we use PLA (Figures 3 and 4) to probe GAPDH-Siah-1 complex under nitrosylation conditions and show that that GAPDH and Siah1 interact under different stimuli of nitrosylation agents. Furthermore, we use R-(-)-Deprenyl – a known inhibitor of GAPDH (*22*) and show that in our cell based system, this molecule abrogates the PLA signal in a dose dependent manner. Finally, our data suggest that PLA can detect and quantify the GAPDH-Siah1 complex; a well-known protein-protein interaction implicated in neurodegeneration (*9-11*) and thus could be a method of choice for probing protein-protein interactions and low abundant protein detection.

**Figure 3.**
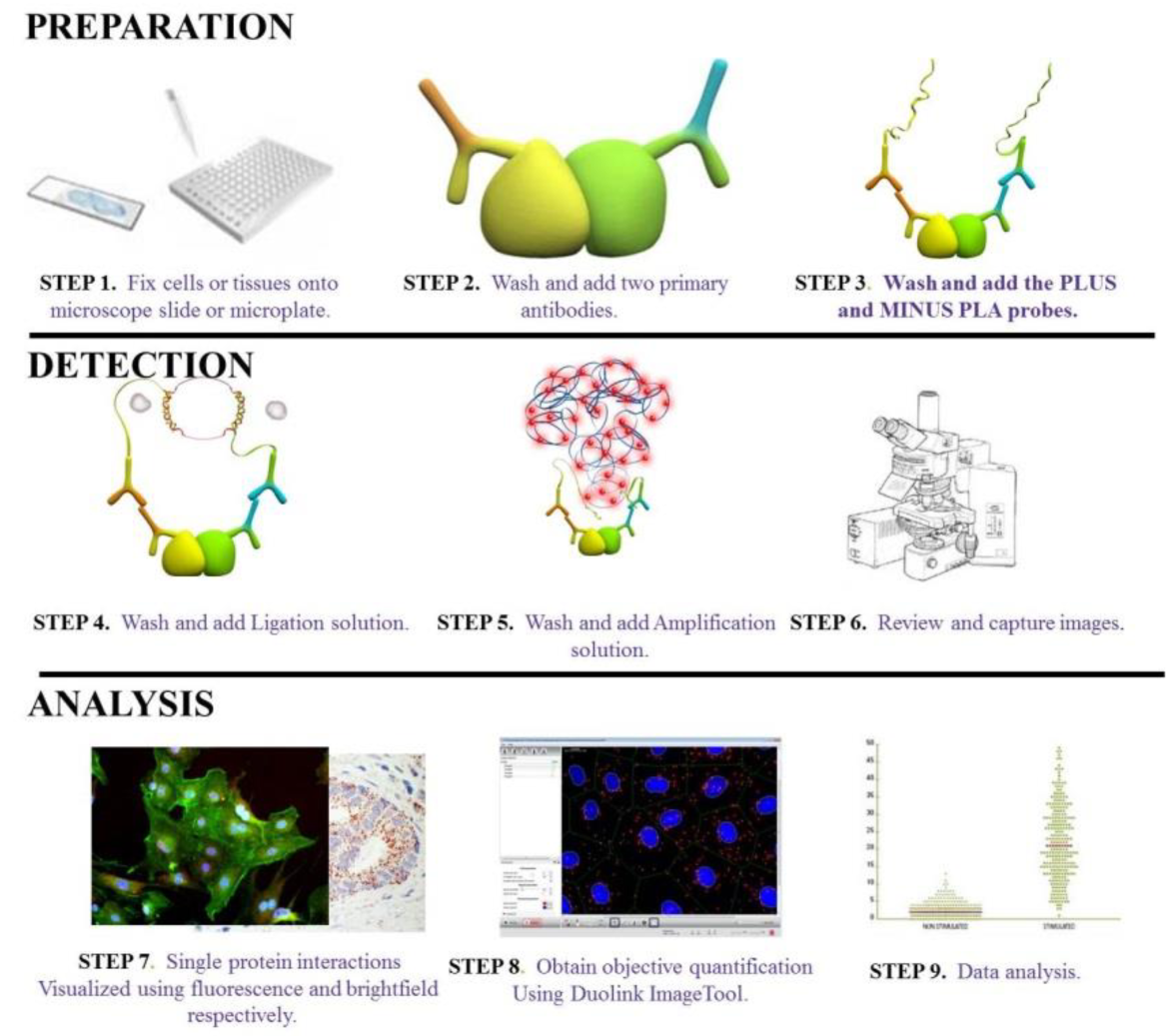
General workflow to probe protein-protein interactions using PLA. Note: Red foci as shown in Steps7 and 8 suggest a PLA signal

**Figure 4.**
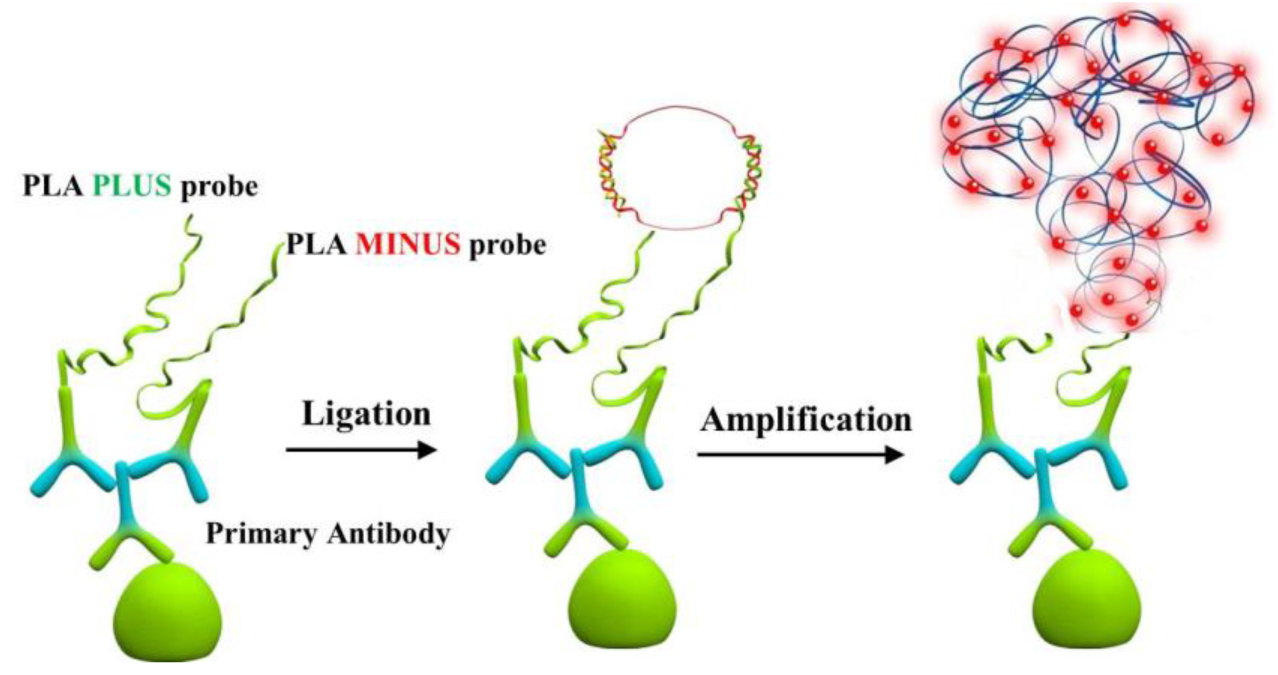
Schematics of protein expression level detection using PLA

## EXPERIMENTAL PROCEDURES

All of the reagents were purchased from MilliporeSigma unless otherwise specifically mentioned.

### Cell culture

Several mammalian cells were investigated in our experiments. Specifically, Human prostate cancer line derived from brain metastasis (DU145); Human gliobastoma multiforme tumor (T98G); Abelson murine leukemia virus-induced tumor (RAW 264.7); Human colon carcinoma cell lines (SW620) and Human breast cancer cell line (MCF-7) were used. All the cell lines were maintained as per the suggested protocols. Briefly, frozen cells were thawed quickly at 37°C and storage media was removed by centrifugation. Resulting cell pellet was resuspended in the appropriate media. Precisely, DU145, T98G, RAW 264.7 and MCF-7 cell lines were propagated into Dulbecco’s Modified Eagle Medium (DMEM)-AQ media, supplemented with 10% (v/v) fetal bovine serum and 1% (v/v) Penicillin-Streptomycin-Glutamine at 37°C and 5% CO_2_. To detach, cell layer was washed with 1X sterile PBS followed by treatment with 0.25% (w/v) Trypsin - 0.53mM EDTA solution and counted using hemocytometer. SW620 cells were cultured using ATCC-formulated Leibovitz’s L-15 Medium supplemented with 10% (v/v) fetal bovine serum and 1% (v/v) Penicillin-Streptomycin-Glutamine at 37°C.

### GAPDH enzyme activity assay

GAPDH enzyme activity assay was performed on DU145 cells treated with 0μM, 500 μM and 1000 μM GSNO respectively. Purified GAPDH was also used as a positive control. Briefly, DU145 cells were grown as mentioned above and pelleted by centrifugation. Cells were res-suspended in 1X PBS pH 7.0 prior to GAPDH enzyme activity assay. Purified GAPDH was dissolved in 250mM Glycylglycine pH 7.4 buffer. 60mM solution of D-Glucose-6-phosphate, 20mM solution of β- nicotinamide adenine dinucleotide phosphate (β-NADP) and 300mM MgCl_2_ solution were prepared separately in deionized water. Next, a cocktail was prepared by mixing 21mL deionized water, 5.0mL of 250mM Glycylglycine pH 7.4 buffer, 1.0mL of 60mM D-Glucose-6-phosphate, 1mL of 20mM β-NADP and 1mL of 300mM MgCl_2_ and pH of the cocktail was checked and adjusted to 7.4 using 1M NaOH solution. 2mM GSNO solution was prepared using 1X PBS pH 7.0. 2.9mL of the cocktail solution was pipetted into a cuvette and incubated at 25°C prior to assay. Assay was started by adding 0.1mL of enzyme solution (GAPDH, GAPDH+1mM GSNO, DU145 Lysate, DU145 Lysate+ 0.5mM GSNO and DU145 Lysate+1mM GSNO). NADPH generation was monitored at 340nm and data was plotted and analyzed.

### Proximity Ligation Assay (PLA)

PLA experiment was performed as per manufacturer’s recommendation. Briefly, a silicon gasket chamber coverslip was coated with fibronectin and incubated for 30 minutes at 37°C. Next, cell confluent at 50-70% level were harvested and plated onto silicon gasket chamber slides. In general, 1000-2000 cells/well were used. This was incubated overnight at 37°C either in the presence or absence of 5% CO_2_ and 90-95% relative humidity. Next day, Nitrosylation reagents (GSNO or SNAP) were prepared in appropriate fresh, room temperature (RT) media and covered with foil. Old media from overnight incubated slides was gently aspirated and freshly prepared GSNO or SNAP was added in different amounts. Specifically, 50μM, 100μM, 150μM, 200μM, 1mM GSNO or 100μM, 200μM, 400μM, 800μM, 1mM SNAP was added to each well. For control experiments, fresh and RT warm media was added. Slides were incubated 24 hours at 37°C either in the presence or absence of 5% CO_2_ and 90-95% relative humidity. Next morning, media was aspirated from slides and 40μL of 4% (v/v) para-formaldehyde solution, prepared in 1X-PBS was added to each well. Slides were incubated at 37°C for 30 minutes and quenched by adding 5μL of 1.25M Glycine prepared in 1X-PBS. This was followed by incubating the slides in permeablization solution (0.75% (v/v) Tween-20 and 0.75% (v/v) Triton X-100) for one hour at RT. Slides were then washed in 1X PBS and 40μL of blocking solution was added to each well and incubated at 37°C for at least one hour. Excess blocking solution was aspirated and primary antibodies, diluted in antibody diluent, were added and incubated overnight at 4°C. PLA certified rabbit anti- GADPH antibody was tested at 10,000 X and 20, 000X dilutions respectively while goat anti- Siah-1 Abcam antibody was tested at 500X, 1000X, 2000X and 4000X dilutions respectively. For a complete PLA reaction between GAPDH and Siah-1, rabbit anti GAPDH antibody was used at 10,000 X dilution and goat anti-Siah1was used at 2,000X dilution. The negative controls for each target were also used at same dilution as described above.

After overnight incubation, antibody solutions were aspirated and slides were washed in Duolink® wash buffer A as per manufacturer’s protocol. Next, a solution consisting 5X dilution of PLA Anti-Rabbit PLUS and Anti-Goat MINUS probes was prepared, 40μL of this solution was added to each well and incubated for 1 hour at RT. Excess solution was aspirated and slides were washed in Duolink® wash buffer A as per manufacturer’s protocol. 40μL of Ligation mix prepared at 40X dilution was added to each well and incubated at 37°C for 30 minutes and washed using Duolink® wash buffer A. 40μL of amplification solution (80X dilution) was added to each well and incubated for 100 minutes at 37°C. Slides were washed in Duolink® wash buffers B and 0.01X wash buffer B respectively as per manufacturer’s protocol. A dye solution containing Phalloidin and Hoechst (both ThermoFisher) was prepared in 1X-PBS and 40μL of which was added to each well at RT for 15 minutes. Slides were washed and results were visualized using Olympus BX51 microscope with DAPI, FITC and CY5 filters for nuclear, actin and Duolink® signal respectively. Images were merged using Olympus cellSens^™^ software.

### Deprenyl Treatment

Deprenyl effect on GAPDH-Siah1 PLA was tested primarily using T98G cell lines due to experimental ease. A fresh 1mM solution of R-(–)-Deprenyl hydrochloride was prepared in cell culture media prior to experiment and stored in dark. To test the effect of Deprenyl, T98G cells were first exposed to 1mM Deprenyl for 30 minutes and after which solution was immediately aspirated and 40μL of Deprenyl at 150μM, 500 μM and 1000 μM respectively supplemented with 150μM GSNO was added to each well. Slides were incubated for 24 hours at 37°C, 5% CO_2_ and 90-95% relative humidity. Cells were next fixed and PLA was performed as described above.

## RESULTS & DISCUSSION

### Cell line validation using PLA

RAW 264.7, T98G and DU145 cell lines were used to establish the suitability of cell lines for PLA. At the outset of experiments, we were interested to investigate the potential translocation of GAPDH to nucleus after its binding to Siah-1. Thus, we conducted experiments to establish cell lines that would allow us to visualize this biological process. We used GAPDH as the target and used single protein detection using PLA to probe the location of GAPDH in cell (Figure 5). This method involves probing the target protein with primary antibody (10,000 X dilution for Anti GAPDH) and then with PLA Anti-rabbit PLUS and Anti-rabbit MINUS probes in next step. The complex thus generated was ligated, amplified and probed using far red detection. Single red foci showed the presence of GAPDH. As expected GAPDH was clearly visible in all three cell lines. However, comparison of PLA data from RAW 264.7 cell lines to that of T98G and DU145 cell lines suggested the use of T98G and DU145 cell lines for probing GAPDH-Siah-1 protein-protein interactions. Specifically, data from RAW 264.7 cell line made GAPDH appear as a nuclear protein under non nitrosylation conditions which is presumably an artifact of RAW 264.7 morphology (Figure 5). On the contrary, data from T98G and DU145 cell lines shows GAPDH as cytosolic protein under identical condition (Figure 5) making it easier for visualizing translocation. Thus, for further experiments, T98G and DU145 cell lines were used.

**Figure 5.**
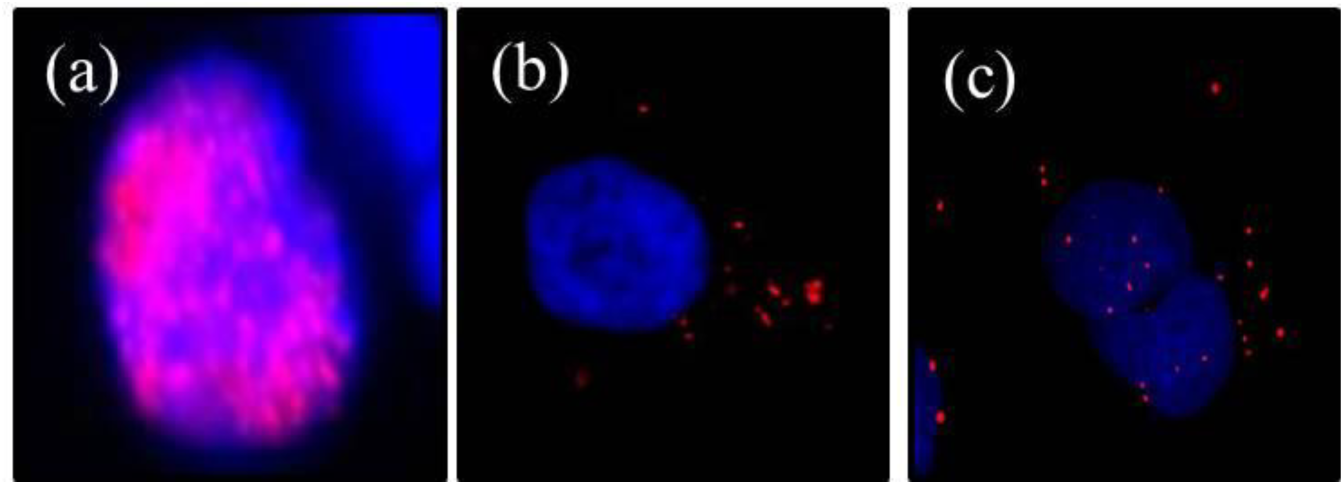
Using GAPDH as target protein, cell line validation via PLA. (a)- Single protein detection using PLA in RAW264.7 cell line, (b)- DU145 cell line, (c)- T98G cell line. Blue-DAPI and Red- CY5 staining

### Antibody validation using PLA

GAPDH and Siah-1 antibodies were validated using western blot and PLA. Our results indicated that western blots were not reproducible (data not shown) when used to probe endogenous levels of Siah-1 protein with various antibodies. We next used PLA to validate antibodies for Siah-1 and GAPDH. For Siah-1, we used single protein recognition PLA as described. Briefly, for Siah-1 we performed PLA at 500X and 1000X anti Siah-1 antibody in RAW 264.7 cells (Figure 6 a-b) and 500X, 1000X, 2000X and 4000X dilution series for DU145 cell lines (Figure 6 c-f). Similarly, for GAPDH antibody validation, we performed PLA at 10,000 X and 20,000X dilutions in DU145 cells (Figure 7 a-b). Based upon our data, we concluded that 2000X Anti-Siah-1 (Goat) antibody and 10,000X Ant-GAPDH (rabbit) gave no background signal in PLA experiment and were suitable dilutions for conducting PLA.

**Figure 6.**
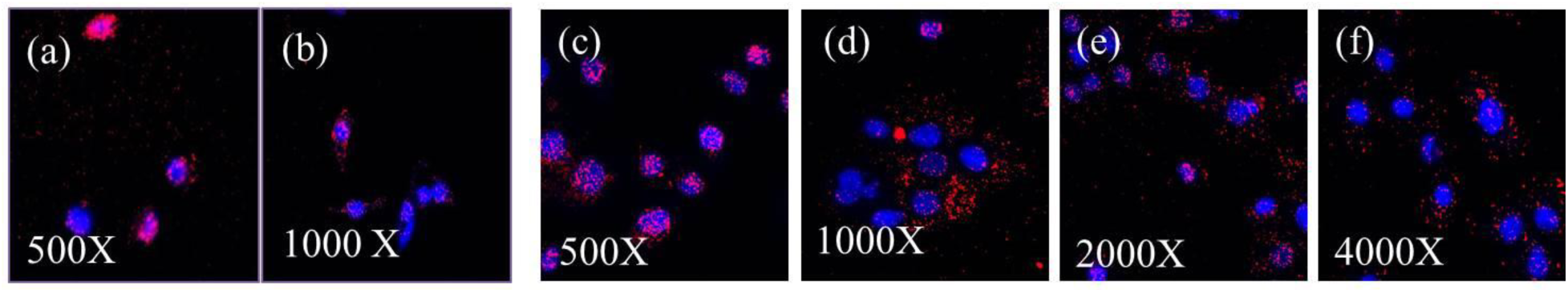
Anti Siah-1 antibody validation using PLA. (a) 500X dilution, (b) 1000X dilution and resulting PLA signal in RAW264.7 cells; (c-f)- 500X, 1000X, 2000X, 4000X dilution and resulting PLA in DU145 cells. Blue- DAPI and Red- CY5 staining.

**Figure 7.**
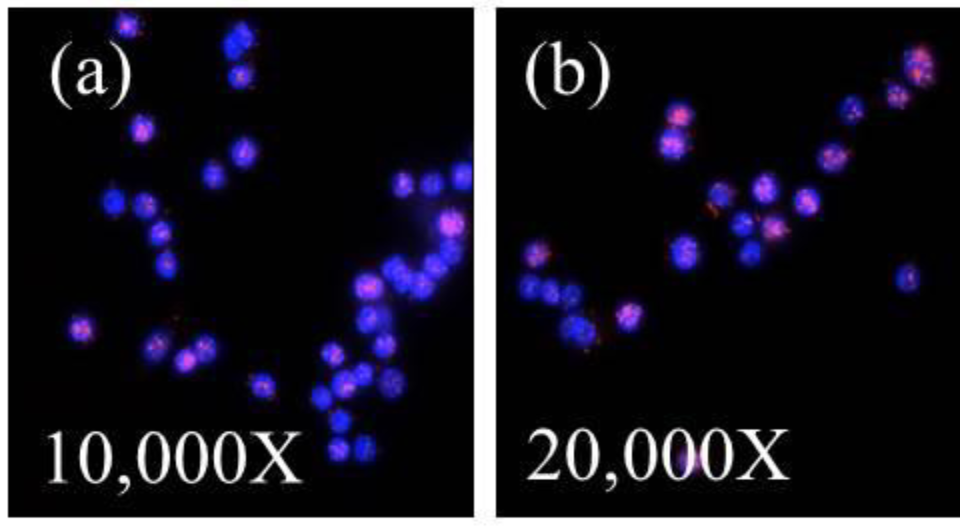
Anti GAPDH antibody validation using PLA. (a)- 10,000 X dilution, (b)- 20,000 dilution and resulting PLA signal in RAW 264.7 cells. Blue- DAPI and Red- CY5 staining.

### GAPDH activity assay

GSNO nitrosylates GAPDH using its active site cysteine and thus abolishes its enzymatic activity(*23*). To verify that in our PLA experiments, GSNO is reducing GAPDH enzymatic activity, we performed GAPDH enzyme assay on DU145 cell lysate in the presence and absence of 0.5mM and 1mM GSNO respectively. Our data shows cell lysate treated with GSNO loses GAPDH enzyme activity over time (Figure 8). For comparison, we also performed the same experiment with purified GAPDH in presence and absence of GSNO and observed similar trends (Figure 9). Reduction of GAPDH enzyme activity in DU145 cells upon treatment with GSNO suggest that GAPDH active site cysteine might be modified as expected and shown by other groups.

**Figure 8.**
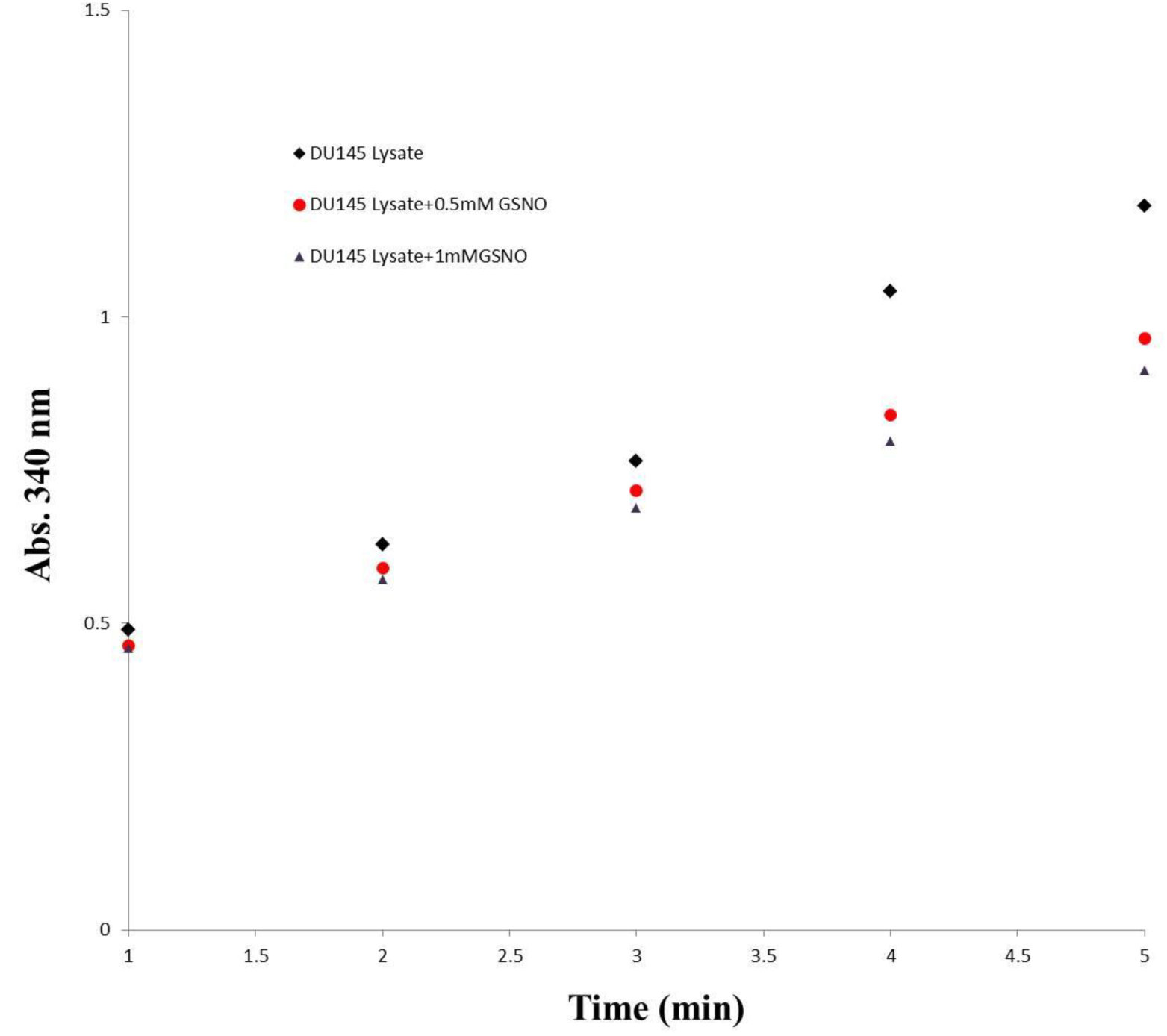
GAPDH activity measurement in DU145 cells in the presence and absence of GSNO

**Figure 9.**
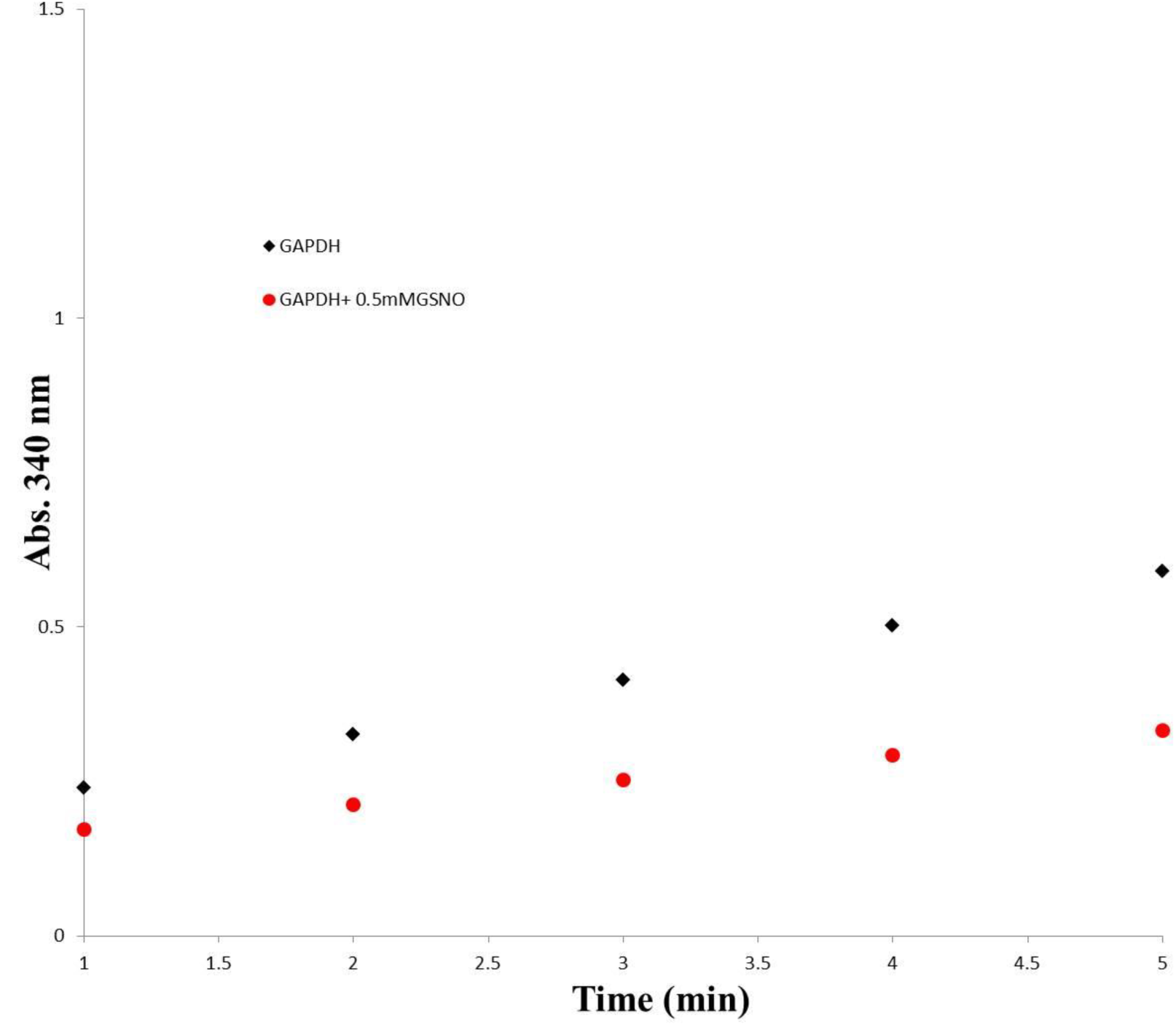
GAPDH activity measurement using purified GAPDH in presence and absence of GSNO

### PLA in the absence of nitrosylation reagents

After antibody validation, we next performed a complete PLA reaction with appropriate technical controls. We first performed PLA experiment between GAPDH and Siah-1 under non-nitrosylation i.e. no GSNO or SNAP conditions. In addition, we also performed three technical controls. In first control, we eliminated Siah-1 antibody; In second, we eliminated GAPDH antibody; and In third, we eliminated both primary antibodies and only used PLA PLUS and PLA MINUS probes. Under non-nitrosylation conditions, our results suggest that GAPDH-Siah-1 show no detectable interactions as visualized using PLA (Figure 10). Data from multiple experiments in DU145 cell line showed the absence of red-foci (PLA signal) in different focal planes. These results validate previous observations and also provide confirmatory visual evidence to the fact that GAPDH nitrosylation may be necessary for its interaction with Siah-1.

**Figure 10.**
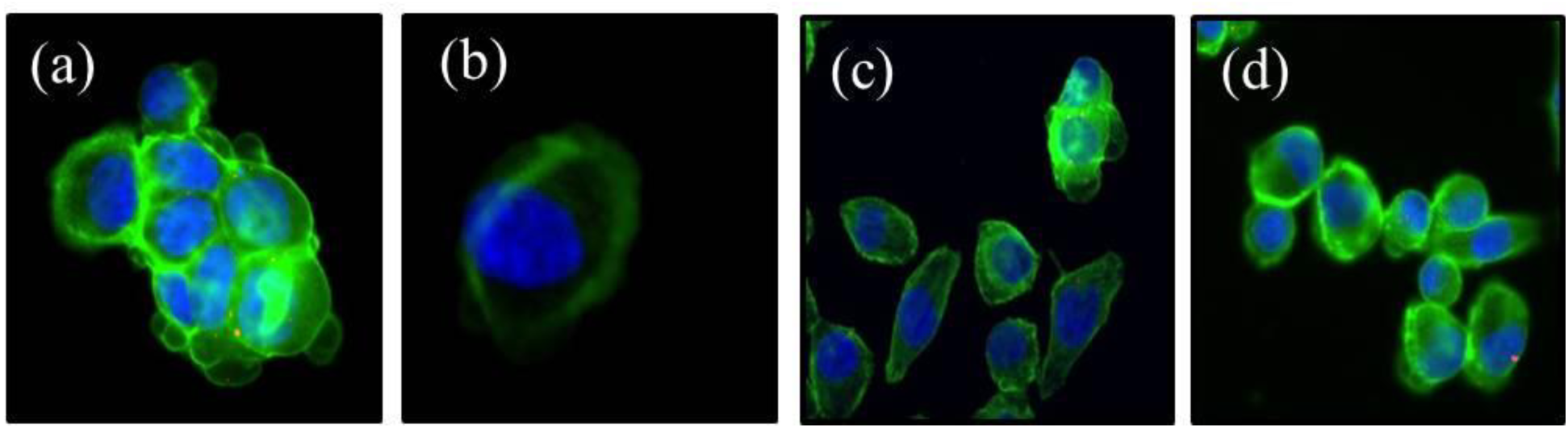
GAPDH-Siah-1 PLA reaction in the absence of nitrosylation agents. (a)- complete PLA reaction in which both (GAPDH and Siah-1) primary antibodies were used, (b)-technical control in which Siah-1 antibody was eliminated, (c)- technical control in which GAPDH antibody was eliminated, (d)- negative control with no primary antibodies. Blue- DAPI; Red-CY5 and Green- Actin staining

### PLA in the presence of nitrosylation reagents

We next performed a complete PLA in the presence of different amounts of GSNO and SNAP reagents respectively. As shown in (Figure 11), we observe a clear PLA signal (red foci) in the complete PLA reaction. Expectedly, no such signal is observed in the technical controls. We obtained consistent positive PLA signal when either of the SNAP or GSNO were used. Our data clearly indicates that GSNO signal seemed more optimal at 150μM concentration while 1000μM SNAP lead to higher PLA signals in DU145 cells (Figures 12 and 13). To verify whether the PLA signal is not cell line dependent, we treated T98G cells with GSNO and SNAP respectively and observed a similar trend (Figure 14). We also quantified the PLA signal between GSNO untreated complete PLA reaction and GSNO treated complete PLA reaction. Our analysis using Duolink^®^ Image Tool suggests that there was a 992% increase (Figure 15) in signal when the cells were treated with GSNO and thus GAPDH nitrosylation might be a critical event for this protein-protein interaction to occur. Nonetheless, our data provides a clear visual evidence of a GAPDH-Siah-1 interaction under the conditions tested.

**Figure 11.**
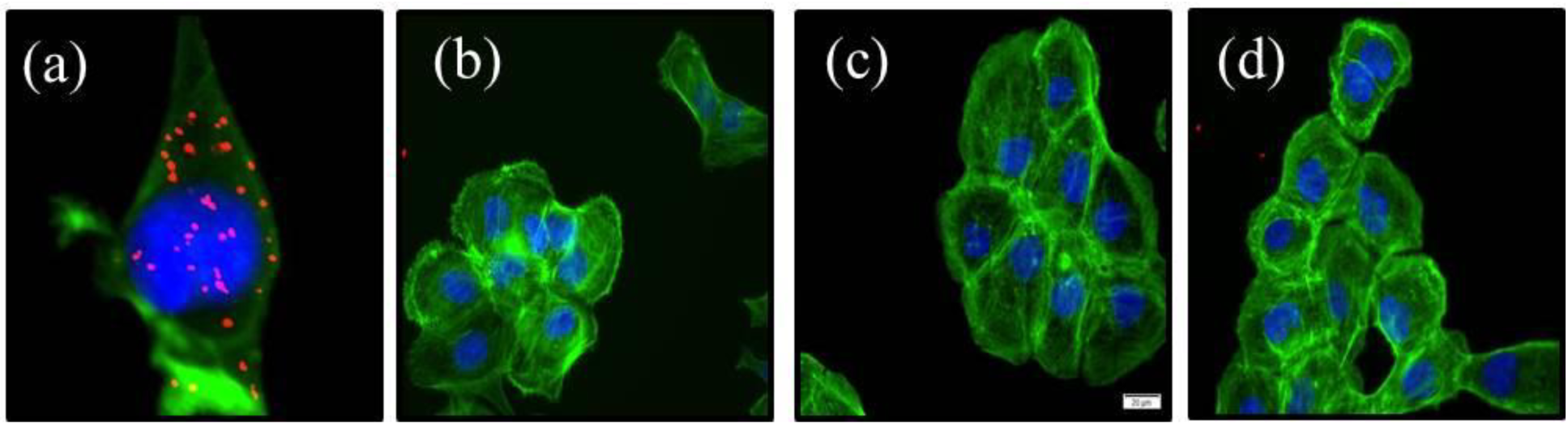
GAPDH-Siah-1 PLA reaction in the presence of 150μM GSNO (a)- complete PLA reaction in which both (GAPDH and Siah-1) primary antibodies were used, (b)-technical control in which Siah-1 antibody was eliminated, (c)- technical control in which GAPDH antibody was eliminated, (d)- negative control with no primary antibodies. Blue- DAPI; Red- CY5 and Green-Actin staining

**Figure 12.**
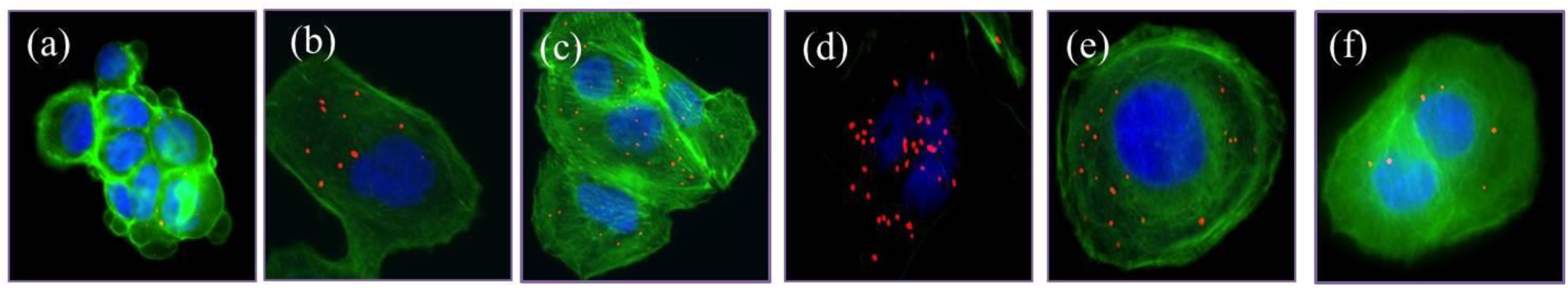
GAPDH and Siah-1 PLA signal response vs. increasing concentration of GSNO in DU145 cells (a)- 0μM GSNO, (b)- 50μM GSNO, (c)- 100μM GSNO, (d)- 150μM GSNO, (e)-200μM GSNO, (f)- 1000 μM GSNO. Blue- DAPI; Red- CY5 and Green- Actin staining

**Figure 13.**
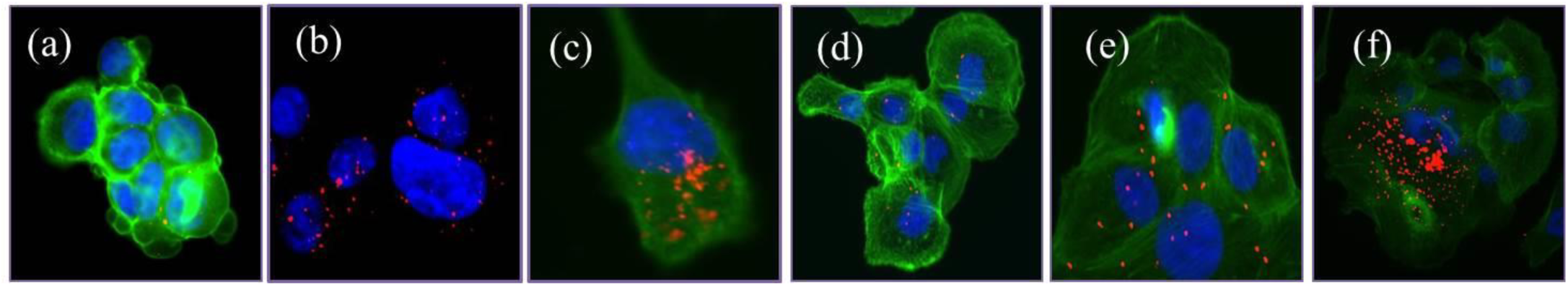
GAPDH and Siah-1 PLA signal response vs. increasing concentration of SNAP in DU145 cells (a)- 0μM SNAP, (b)- 100μM SNAP, (c)- 200μM SNAP, (d)- 400μM SNAP, (e)-800μM SNAP, (f)- 1000 μM SNAP. Blue- DAPI; Red- CY5 and Green- Actin staining

**Figure 14.**
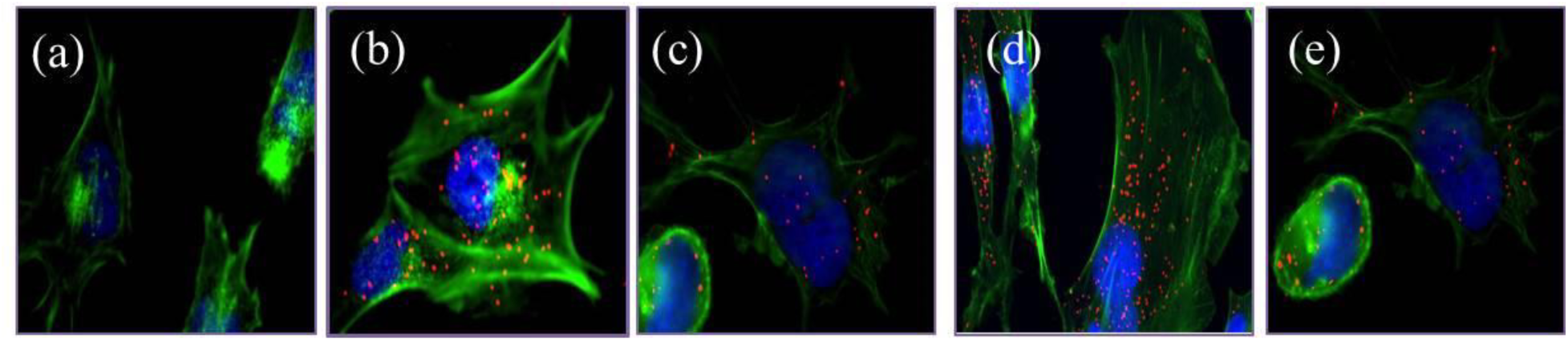
GAPDH and Siah-1 PLA signal response vs. increasing concentration of GSNO and SNAP in T98G cells (a)- no nitrosylation agent, (b)- 150μM GSNO, (c)- 1000μM GSNO, (d)-200μM SNAP, (e)- 1000 μM SNAP. Blue- DAPI; Red- CY5 and Green- Actin staining

**Figure 15.**
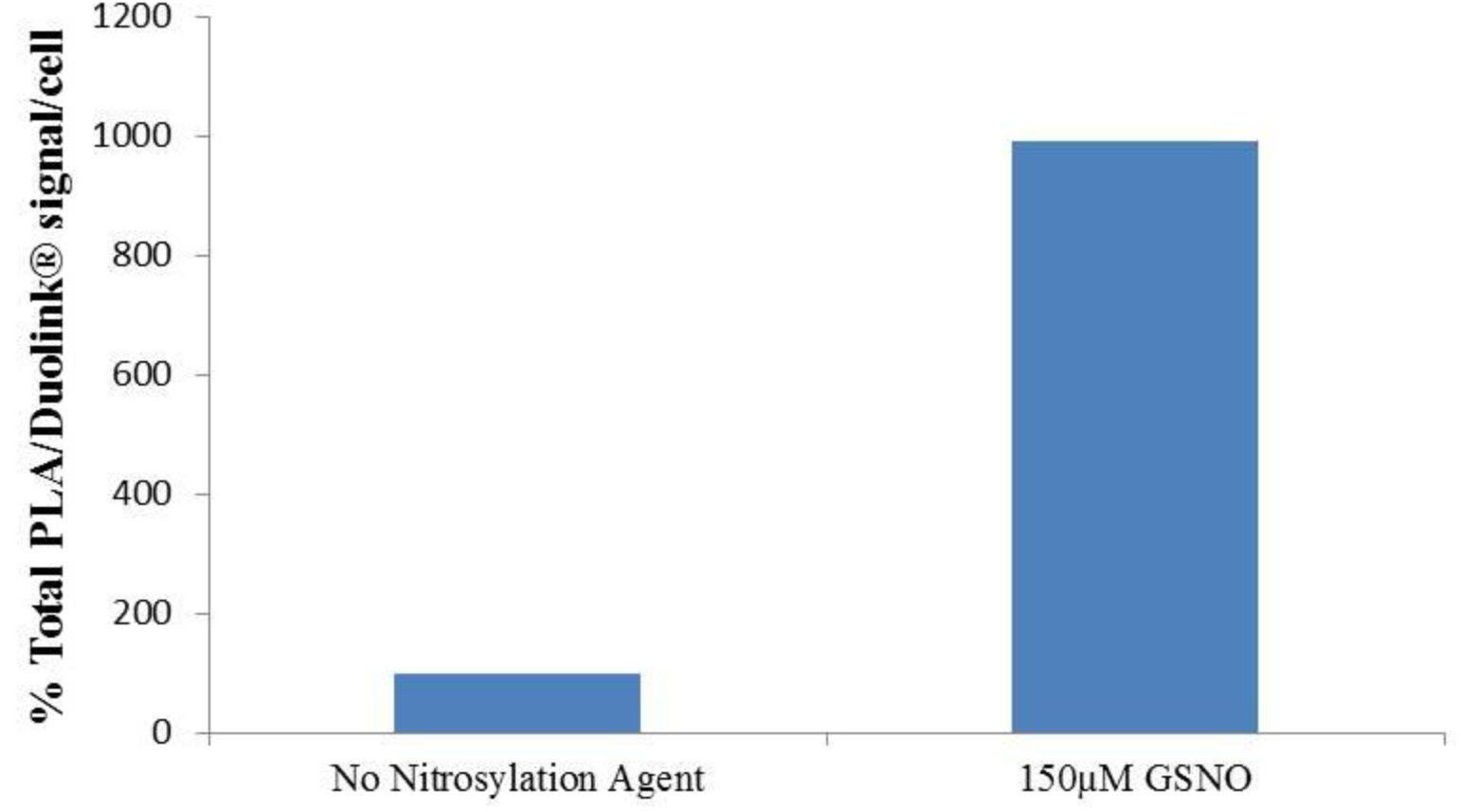
Quantification of PLA/Duolink® signal shows 990% increase in signal in GSNO untreated vs GSNO treated (150μM) T98G cells.

### Inhibition of GAPDH-Siah-1 PLA using Deprenyl

GAPDH is a critical housekeeping gene with significant implications in nuclear events and apoptosis. GAPDH-Siah-1interactions, provoked by cellular stress such as nitrosylation, has been shown to be involved in Parkinson’s disease, AD (*19*), cerebral ischemia–reperfusion injury(*20*) and acute lung injury(*21*). Thus we decided to use our PLA based cell assay to investigate the effect of deprenyl, which has been shown to inhibit GAPDH and act as potential drug candidates for PD and other neurodegenerative disorders. Our results suggest that 150μM deprenyl for 24 hours reduce GAPDH-Siah-1 PLA signal by ∼70% (Figure 16). Interestingly, *Hara et al*.(*11*) showed that GAPDH preincubated with deprenyl at 0.01nM and 1nM respectively, significant inhibits GAPDH-Siah-1 association in a co-immunoprecipitation experiment. We note that the significantly higher amounts of deprenyl were needed in our experiments to observe GAPDH-Siah1 complex inhibition. One reason for this could be that our experiments are conducted in cells where one might envision a crowded environment requiring higher amounts of deprenyl to achieve significant inhibition. Additionally, it is a well-known that deprenyl is also an inhibitor for other proteins such as monoamine oxidase-B (MAO-B) (*24, 25*) and thus an off target binding of deprenyl in crowded environment could explain our results. Nonetheless, our data not only provides confirmatory but visual evidence of GAPDH-Siah-1 binding but also a cell based model system that could be used to investigate anti-apoptotic and anti-neurodegenerative compounds in biologically relevant settings.

**Figure 16.**
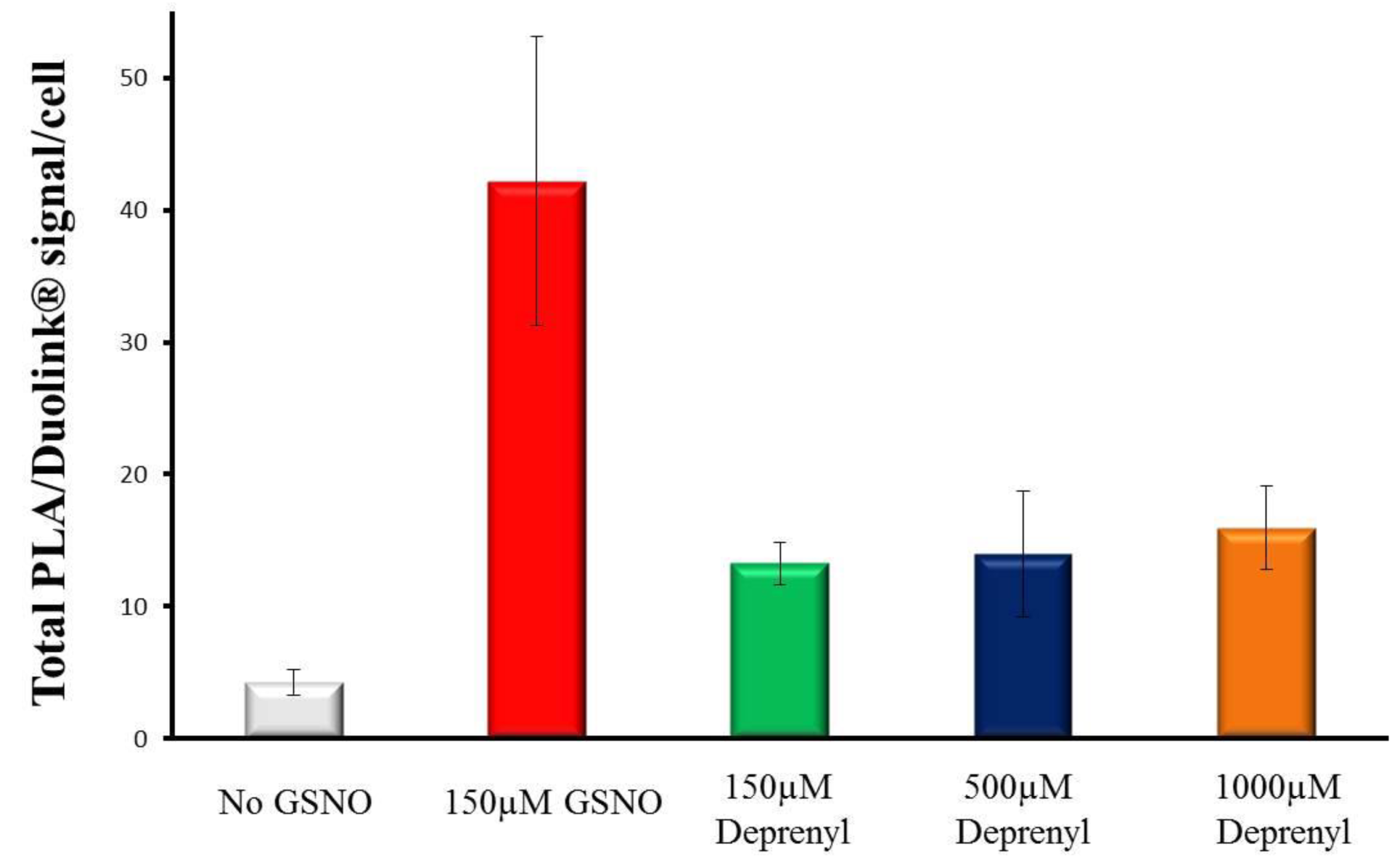
Quantification of PLA/Duolink^®^ signal in the absence and presence of Deprenyl in T98G cells.

## Author Contributions

H.S. conceived the project and designed research. H.S. and C.M. conducted experiments. H.S. and C.M. analyzed the data. H.S. wrote the manuscript.

## Acknowledgements

Authors would like to thank Dr. Vikas Palhan for help with microscopy, Mr. Adam Kronk for help with initial reading of the draft, Mr. Thomas Juehne and Dr. Jeffrey Turner for critical suggestions.

